# Behavioral plasticity and the valence of indirect interactions

**DOI:** 10.1101/2024.08.15.608151

**Authors:** Ashkaan K. Fahimipour, Michael A. Gil, Andrew M. Hein

**Affiliations:** Department of Biological Sciences, Florida Atlantic University, Boca Raton, FL, USA; Center for Complex Systems, Florida Atlantic University, Boca Raton, FL, USA; Department of Ecology and Evolutionary Biology, University of Colorado, Boulder, CO, USA; Department of Computational Biology, Cornell University, Ithaca, NY, USA

**Keywords:** predator-prey, coarse-graining, competition, mutualism

## Abstract

Behavioral plasticity in animals influences direct species interactions, but its effects can also spread unpredictably through ecological networks, creating indirect interactions that are difficult to anticipate. We use coarse-grained models to investigate how changes in species behavior shape these indirect interactions and influence the broader dynamics of ecological networks. As an illustrative example, we examine predators that feed on two types of prey, each of which temporarily reduces activity after evading an attack, thereby lowering vulnerability at the expense of growth. We demonstrate that this routine behavior can shift the indirect interaction between prey species from apparent competition to mutualism or parasitism. These shifts occur when predator capture efficiency drops below a critical threshold, causing frequent hunting failures. As a result, one prey species indirectly promotes the growth of the other by relaxing its density dependence through a cascade of network effects, paradoxically increasing predator biomass despite decreased hunting success. Empirical capture probabilities often fall within the range where such dynamics are predicted. We characterize these shifts in the qualitative nature of species interactions as changes in *interaction valence*, highlighting how routine animal behaviors reshape community structure through cascading changes within ecological networks.

## INTRODUCTION

Instances where seemingly entrenched interactions between species abruptly change have now been observed in most ecosystems and across taxa. Included are shifts from competition to facilitation among terrestrial plants in harsh environments (Bertness & Callaway 1994) or bacteria in biofilms (Nadell *et al*. 2016); dynamic shifts between competition and predation (Fahimipour & Anderson 2015), competition and mutualism (M. A. Gil & Hein 2017), or parasitism and mutualism in fishes (Bshary *et al*. 2008); the induction of cannibalism among competing protozoans (Giese 1938), rotifers (Gilbert 1973), and amphibians (Collins & Cheek 1983); or even reversals of predator and prey identity among marine invertebrates (Barkai & McQuaid 1988).

These examples emphasize how changes in the behavior or physiology of interacting organisms can alter the *valence* — the qualitative nature — of their interactions (Huang & Sih 1990; Winnie Jr 2012; Fahimipour & Anderson 2015; M. Gil *et al*. 2019; Gaiarsa *et al*. 2021). Such shifts can significantly impact ecosystem dynamics (Neutel *et al*. 2007) through indirect effects that propagate through ecological networks (Dunne & Williams 2009; Fahimipour, Anderson & Williams 2017). For instance, predators not only reduce prey numbers through consumption but also induce behavioral changes that affect prey demographics and interactions with other species (Schmitz, Beckerman, *et al*. 1997; Schmitz, Krivan, *et al*. 2004). This is evident in behaviorally mediated trophic cascades (Schmitz, Beckerman, *et al*. 1997; Schmitz, Krivan, *et al*. 2004), where prey’s fear of depredation increases biomass in lower trophic levels (Abrams *et al*. 1996; Beckerman *et al*. 1997; Schmitz, Beckerman, *et al*. 1997; Schmitz & Suttle 2001; Schmitz, Krivan, *et al*. 2004). However, our understanding of how behavior influences other indirect effects, including inherently indirect interactions like apparent competition (Holt 1977; Holt & Bonsall 2017; Schreiber & Křivan 2020), remains less developed (Schreiber, Bürger, *et al*. 2011).

We present a theoretical approach for understanding how behavioral plasticity shapes indirect interactions in ecological networks. A central component of this approach is coarse-graining animal behavior into discrete functional states (Fig. 1), which allows us to systematically incorporate fast within-individual behavioral shifts into slower population dynamics. By integrating behavioral coarse-graining with analyses of press perturbations in networks (Bender *et al*. 1984; Nakajima 1992; Yodzis 2000; Aufderheide *et al*. 2013), we reveal how behavioral changes propagate through ecological communities in unexpected ways. We illustrate this phenomenon using a well-studied motif: a predator feeding on two prey species that do not directly compete, a scenario known as apparent competition (Holt 1977; Holt & Bonsall 2017; Schreiber & Křivan 2020). As an example, we examine how a widespread antipredator strategy — hiding, where prey temporarily reduce foraging activity and hence vulnerability in response to predator encounters (Sih 1987; Lima & Dill 1990; Sih *et al*. 2004; Heithaus *et al*. 2009; Herberholz & Marquart 2012) — modifies indirect effects between apparent competitors. By coarsening behavior into transitions between two states (*hiding* or *not hiding*), we show that prey can shift between apparent competition, mutualism, or even parasitism-like relationships, depending on ecological parameters.

**Figure 1.**
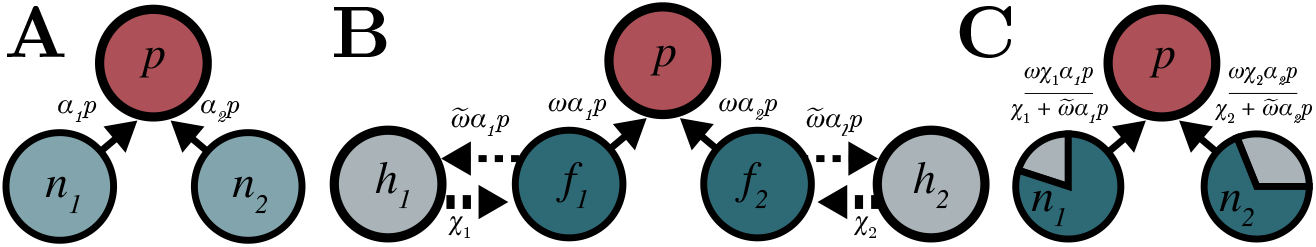
Model coarse-graining approach. Networks illustrate biomass flows in the system. Solid arrows denote trophic links, while dashed lines represent biomass flows due to behavioral state transitions. The *per capita* rates of each reaction are indicated, with variables and parameters in Table 1. **A**. Coarse-graining begins by defining a conventional ecological model. **B**. Model is expanded by subdividing one or more species into functionally distinct subpopulations. **C**. Using separation of time scales, the model is reduced back to its original dimensionality, with newly derived functions governing interactions.

## METHODS

Our approach integrates two established methods: model coarse-graining through time scale separation (Michaelis, Menten, *et al*. 1913; Holling 1959; Real 1977; Auger & Roussarie 1994; Dawes & Souza 2013) and analysis of press perturbation impacts (Bender *et al*. 1984; Nakajima 1992; Yodzis 2000; Aufderheide *et al*. 2013; Novak *et al*. 2016) to reveal how animal behaviors reshape interactions among indirectly connected species. We illustrate this approach by starting with a conventional model of apparent competition (Fig. 1A) and then expanding it to partition populations into distinct behavioral subpopulations (Fig. 1B). This expansion allows us to derive a reduced model that retains the original dimensionality (Fig. 1C) while capturing the population-level consequences of behavioral dynamics. Although we focus here on a specific network motif and behavior, our method can be readily extended to other functional shifts in populations or to *n*-species food webs with only minor modifications.

**Table 1:**
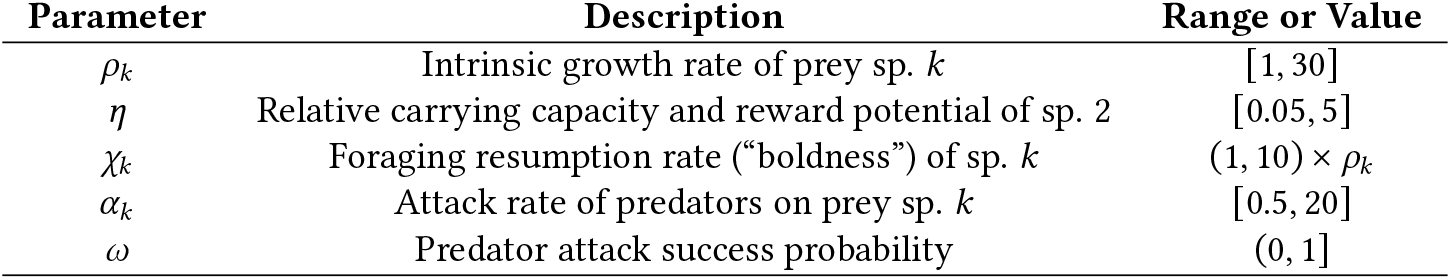
Parameter definitions and ranges used in sensitivity analysis of interaction valences.

### AN EXPANDED MODEL OF APPARENT COMPETITION

The apparent competition motif consists of two prey species that are consumed by a shared predator (Holt 1977). Suppose that we divide both prey species *k* ∈ [1, 2] into two sub-populations each: the “foragers” (denoted as *F*_*k*_), actively foraging in the local habitat, and the “hiders” (denoted as *H*_*k*_), which have entered a state of reduced activity following a failed predation attempt (Fig. 1). Assuming encounters between species are governed by mass-action, we have

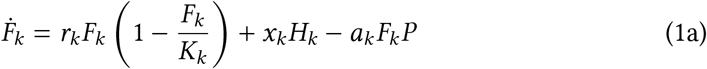

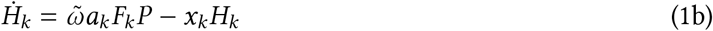

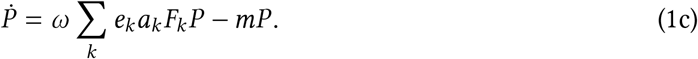

Here, *F*_*k*_ are foraging prey; *H*_*k*_ are prey that are hiding; and *P* is a shared top predator. The parameters *r*_*k*_ are prey growth rates (t^−1^); *K*_*k*_ are prey carrying capacities (individuals · area^−1^); *x*_*k*_ are rates at which hiding prey resume foraging (t^−1^); *a*_*k*_ are attack rates of predators on prey (area · predator^−1^ · t^−1^) that dictate total consumption; *ω* and 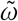 are predator probabilities of attack capture success and failure respectively, with 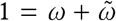; *e*_*k*_ are trophic efficiencies (predator · prey^−1^); and *m* is the predator mortality rate (t^−1^). We assume: [*i*] prey are temporarily safe from predators while in the transient hiding state; [*ii*] they acquire only enough resources to sustain their metabolism, with negligible growth and reproduction (Sih *et al*. 2004); [*iii*] their limited resource acquisition means they contribute very little to density dependence (*i*.*e*., the population’s distance from carrying capacity); [*iv*] prey that escape predators always hide; and [*v*] hiding does not involve competition for finite shelters or habitats, allowing an arbitrary number of prey to hide at once. Assumptions *ii* and *iii* in particular correspond to cases in which prey hiding durations are relatively short and growth is resource-limited (Martín, López & Cooper Jr 2003; Martín & López 2015). In *Appendix S1* we demonstrate that our results are robust to relaxing these assumptions.

### NONDIMENSIONALIZATION

To make the model more generalizable and reduce both the number of parameters and sensitivity to specific parameter values, we first transform variables and parameters into nondimensional quantities (Gurney & Nisbet 1998; Hastings & L. J. Gross 2012). This nondimensionalization step allows us to explore a wide range of ecological scenarios without being limited to specific parameter values, focusing instead on the relative importance of each parameter across orders of magnitude (Table 1). The result is a model in which we have rescaled rates by the mean generation time of top predator individuals (*m*^−1^). Making the following substitutions:

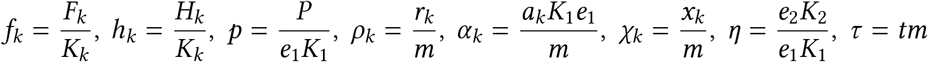

we have the nondimensionalized system of equations (Fig. 1)

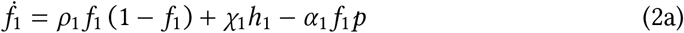

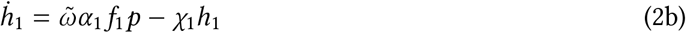

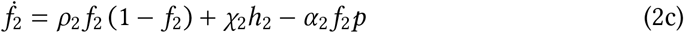

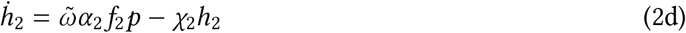

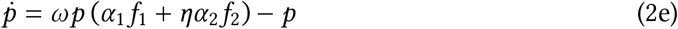

where the derivatives are with respect to our new units of time. Time scale separation

We assume that prey behavioral transitions are (at least approximately) fast compared to prey population demographics (*i*.*e*., prey transition in and out of the hiding state many times during their lives), and that both of these processes are faster than predator mortality, implying 1 < *ρ*_*k*_ < *χ*_*k*_. This allows us to reduce the system using a separation of time scales. For total prey biomass *n*_*k*_ = *f*_*k*_ + *h*_*k*_, we make the substitution *f*_*k*_ = *n*_*k*_ − *h*_*k*_ and differentiate to obtain the slow dynamics

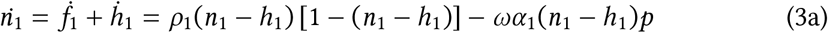

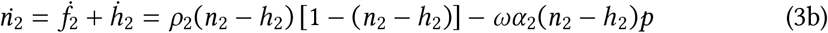

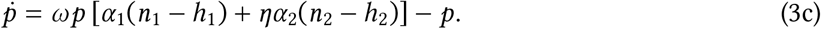

We now consider the fast dynamics of *h*_*k*_. Assuming these fast variables quickly reach their steady state values 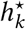, we set 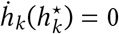 (Eq. 2) to find 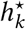:

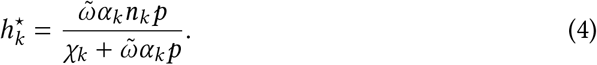

Substituting the steady state densities of hiding individuals 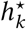 back into the equations for 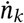, and 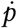 (Eq. 3), we obtain the final reduced system (Table 1)

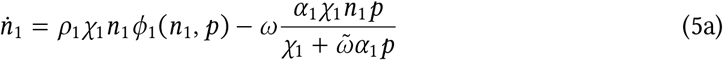

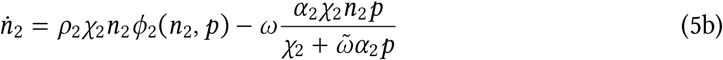

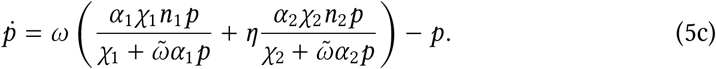

We refer to the normalized rate at which hiding prey resume foraging, *χ*_*k*_, as species “boldness” since higher values of this parameter indicate quicker returns to foraging after successfully escaping a predator attack. The function *ϕ*_*k*_(*n*_*k*_, *p*) is

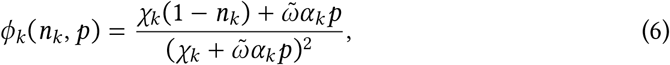

which defines density dependence and the effects of hiding on prey population growth, representing three processes. For prey spp. *k*, in addition to classical self-regulation, *χ*_*k*_(1 − *n*_*k*_), the growth function *ϕ*_*k*_(*n*_*k*_, *p*) captures the dual nature of hiding in our model. Predator-induced hiding provides temporary safety for prey individuals and weakens density dependence at the population scale, enhancing growth 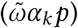, but it also causes lost foraging opportunities, leading to factors 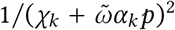. This balance between linear growth-enhancing and quadratic growth-suppressing effects of hiding (and hence, predator densities, which drive prey hiding behavior) is crucial in shaping the interaction dynamics. We note that the model reduces to standard apparent competition with logistic prey growth and type-I functional responses when predators are perfectly efficient hunters, *ω* = 1; 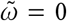. In this case, hiding is never induced and so its effects on population demographics are not realized.

### ANALYSIS OF INTERACTION VALENCE

The local stability of steady states 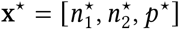 can be computed from linearizations of the dynamics, captured by the Jacobian matrix **J**, which represents dynamics of the system near its equilibrium. The entries of **J** capture the local sensitivities of each variable to small perturbations,

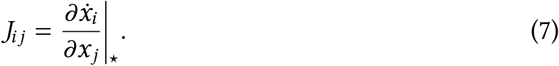

Our initial focus lies in understanding coexistence patterns and hence the location of transcritical bifurcations (Kuznetsov 1998) in the reduced model. We derive analytical test functions (T. Gross & Feudel 2004) to locate these bifurcations in parameter space, which occur when the determinant of the Jacobian matrix vanishes, |**J**| = **0**, marking the onset of a zero eigenvalue (Kuznetsov 1998).

Our next goal is to understand how the interaction between prey species, and between predator and prey, changes with model parameters. To do this we quantify the net interaction between species by accounting for direct and indirect effects through all paths in the motif (Bender *et al*. 1984; Nakajima 1992; Yodzis 2000; Aufderheide *et al*. 2013; Novak *et al*. 2016), which can be accessed directly from the Jacobian. Consider conceptually that we compute the steady state of the system as a function of parameters *s*, which we can write as x^⋆^ = *x*^⋆^(*s*). Since we define our system as 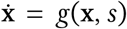 we know that *g*(x^⋆^(*s*), *s*) = 0. Using a corollary to the implicit function theorem (Khalil 2002; Aufderheide *et al*. 2013) we can differentiate with respect to *s* to get

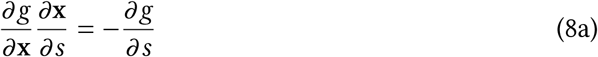

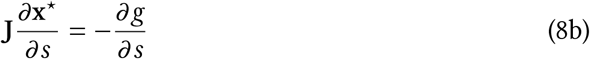

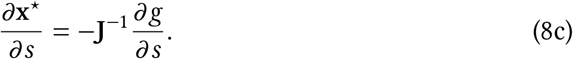

Thus, the impact matrix that captures direct and indirect effects among species is **M** = −**J**^−1^ (discussed in detail by Bender *et al*. 1984; Nakajima 1992; Yodzis 2000; Aufderheide *et al*. 2013; Novak *et al*. 2016), where the superscript indicates a matrix inverse. While we analyze the full matrix, the off-diagonal entries *M*_1,2_ and *M*_2,1_ are particularly informative since they permit a straightforward prey interaction classification: *apparent competition* occurs when both prey negatively affect each other, max(*M*_1,2_, *M*_2,1_) < 0; *apparent parasitism* occurs when one prey benefits at the expense of the other, min(*M*_1,2_, *M*_2,1_) < 0 < max(*M*_1,2_, *M*_2,1_); and *apparent mutualism* occurs when both prey benefit from one another, 0 < min(*M*_1,2_, *M*_2,1_). This classification relies on the linear approximation of Eq. 8, which remains valid for small biomass perturbations.

To assess the tendency of the system to exhibit any of the indirect interaction modes outlined above, it is important to account for the effects of parameter variation in Eq. 5. By parameter variation, we mean the process of randomly drawing parameter values from biologically relevant distributions (Eq. 5, Table 1), which can influence whether the system exhibits certain dynamics. We quantify this tendency as the robustness, ℛ, of a given interaction type to variation in the parameter space of Eq. 5 (Lawton *et al*. 2024). Specifically, we define a binary variable Ω_*q*_(*s*), which equals 1 if the *n*-dimensional vector of model parameters *s* corresponds to interaction type *q*, and 0 otherwise. The fraction of parameter space resulting in interaction type *q* is approximated over *N* = 10^6^ random parameter set samples *s*_*i*_ (Table 1) as

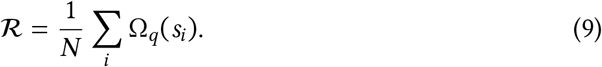

This quantity approximates the region of parameter space where all species coexist 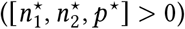 and the indirect prey interaction takes a specific form, providing a measure of each interaction type’s robustness to parameter variation. Our focus on nondimensionalized models reduces the dependence on specific, absolute parameter values and focuses instead on the *relative* influence of different parameters on system behavior (Gurney & Nisbet 1998; Hastings & L. J. Gross 2012). We therefore sweep parameters over multiple orders of magnitude, ensuring a broad exploration of possible ecological regimes.

We performed a logistic regression to quantify how the presence of apparent mutualism depends on the set of parameter values (Table 1). More specifically, we used a generalized linear model with a binomial error distribution and a logit link function. For each random parameter set, we defined a binary variable indicating the presence or absence of mutualism, where mutualism was defined as min(*M*_1,2_, *M*_2,1_) > 0. We note that the linear model was not used to assess statistical significance (White *et al*. 2014), but rather to quantify how parameters influence the probability of mutualism.

## RESULTS

We begin by analyzing model steady states, focusing on a scenario where the two prey species differ in their hiding strategy as defined by boldness, *χ*_*k*_ — the rate at which hiding individuals of sp. *k* resume foraging — but are otherwise identical in terms of intrinsic growth rates (*ρ*_1_ = *ρ*_2_), carrying capacities, and reward potentials (*η* = 1). Under these conditions, we investigate the effects of changing predator preference for either prey (relative values of *α*_1_ and *α*_2_) and the capture success probability of predators, *ω* (see Eq. 5) on equilibrium biomass x^⋆^. Most of the assumptions made here will be relaxed in the following section, where we present sensitivities of interaction patterns to parameter variation.

### STEADY STATES AND INTERACTION MODES

Assume prey species 2 is bolder than species 1 (*χ*_1_ < *χ*_2_), meaning individuals of species 2 spend less time hiding after evading an attack on average. We consider two distinct cases. First, when *α*_1_ > *α*_2_, predators prefer the shy prey species 1. This could occur if some inherent preference for species 1 causes longer hiding times in that species as an adaptive response (*e*.*g*., Holt & Bonsall 2017; Lima 1998). Alternatively, when *α*_1_ < *α*_2_, the predator prefers the bolder prey species 2. This might happen if the boldness of prey species 2 corresponds with some other behaviors or characteristics that make them easier to encounter or detect (*e*.*g*., Abrams 1984; Werner *et al*. 1983). This preference may also occur in cases where the predator encounters the bolder species more often, allowing it to develop better search images or attack strategies (Endler 1988).

Exploring the model steady states numerically, we find that interaction outcomes and biomasses depend strongly on predator capture probability *ω* and its relative preference for either the bold (*α*_1_ < *α*_2_) or shy (*α*_1_ > *α*_2_) prey (Fig. 2A-C). Across a range of prey preferences, equilibrium biomass is maximized for all species, including predators, at *low* values of predator capture probability, *ω*, (Fig. 2A-C) near the transcritical bifurcation where predators are lost (Fig. 2D, bottommost black curve). This pattern holds across different preferences for either prey (*i*.*e*., relative values of *α*_1_ and *α*_2_). This result is intuitive for prey species, which benefit from reduced top-down pressure as their shared predator becomes an inefficient hunter. However, it is surprising with respect to the predator species, which experiences fitness benefits as its attacks become less likely to succeed (Fig. 2C).

**Figure 2.**
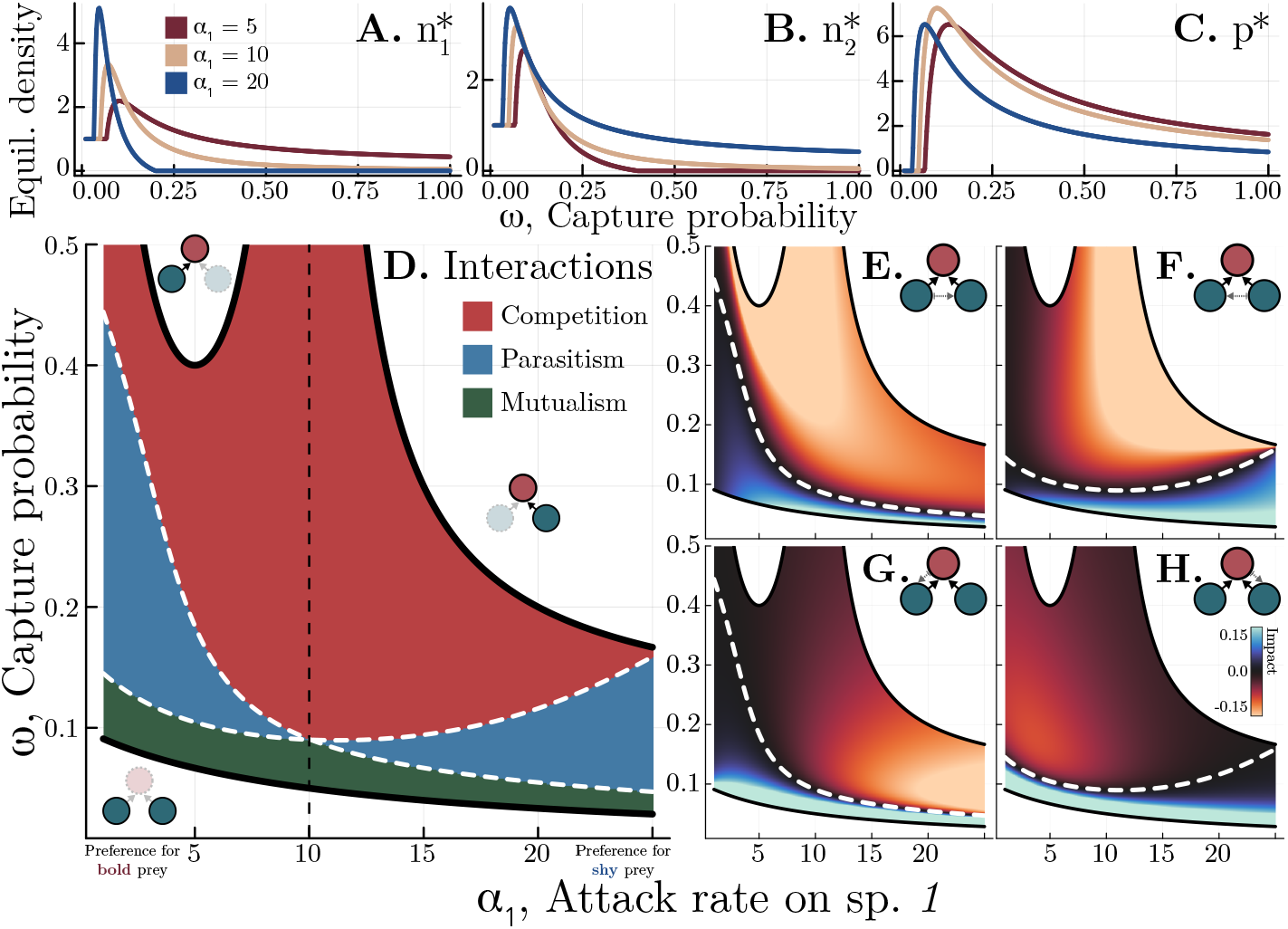
Steady states and interaction valence. **A**. Equilibrium biomass densities of prey species 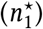, **B**. sp. 2 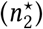, and **C**. the predator (*p*^⋆^) as predator capture probability (*ω*) and attack rate on sp. 1 (*α*_1_) vary. **D**. Interaction valences. Black bifurcation curves mark extinction boundaries, with inset networks showing species losses. White curves mark transitions between interaction modes, *M*_*i,j*_ = 0 (Eq. 8). Vertical line shows the attack rate on the bolder prey species 2 (*α*_2_) for reference. **E**. Impacts of prey sp. 1 on sp. 2, **F**. sp. 2 on sp. 1, **G**. predator on sp. 1, **H**. predator on sp. 2. Parameters are *ρ*_*k*_ = 14.5, *α*_2_ = 10, *η* = 1, *χ*_1_ = 15, *χ*_2_ = 16.

To explain the counterintuitive increase in predator biomass despite decreasing hunting success, we analyzed the full suite of interactions within the motif. Our results reveal a sharp threshold in predator capture probability, *ω*, below which the interactions among species shift from negative (apparent competition) to positive (apparent mutualism or parasitism; Fig. 2D-H). This shift arises from two key effects of reduced hunting success. The first involves lowered prey mortality and weakened density dependence. With lower capture success, fewer prey are killed, and larger proportions cycle in and out of the hiding state. Hiding not only temporarily protects prey individuals from immediate predation but also reduces their contribution to density-dependent population regulation (Eq. 6; Appendix S1). As hiding becomes more common, the direct negative impact of predation is increasingly offset by the benefits of reduced intraspecific competition (Fig. 2G-H). This eventually leads to cascading positive effects among prey: the rise in predator biomass stemming from the above process further induces hiding in the alternative prey species.

This additional shift enhances the reduction in density dependence, ultimately causing the indirect interaction between prey to transition from negative to positive (Fig. 2D–F). In effect, the predator relaxes density dependence in both prey populations, so that each one facilitates the growth of the other through a positive feedback.

Predator biomass is maximized near the critical *ω* where the white curves in Fig. 2D (indicating vanishing impacts of each prey on the other) intersect (compare Figs. 2C and 2D). At this point, the prey populations become effectively decoupled, and the predator exploits two independent resources leading to an equilibrium that resembles apparent neutrality.

### ROBUSTNESS OF APPARENT MUTUALISM

The likelihood of apparent mutualism between prey depends on model parameters. Fig. 3A shows how parameter values correlate with the probability that a randomly-constructed stable community described by Eq. 5 exhibits apparent mutualism. Negative coefficients indicate that higher values of a parameter reduce the odds of mutualism. Apparent mutualism is most common in communities with inefficient predators (small *ω*; Fig. 2D; Fig. 3), longer prey hiding times (small *χ*_*k*_), weaker consumer interaction strengths (small *α*_*k*_), faster intrinsic prey growth (large *ρ*_*k*_), and greater asymmetry in prey rewards or carrying capacities (small *η*). Surprisingly, prey traits such as intrinsic growth rates (*ρ*_*k*_) and hiding latencies (*χ*_*k*_) have the weakest influence on mutualism probability, while predator properties play a larger role.

**Figure 3.**
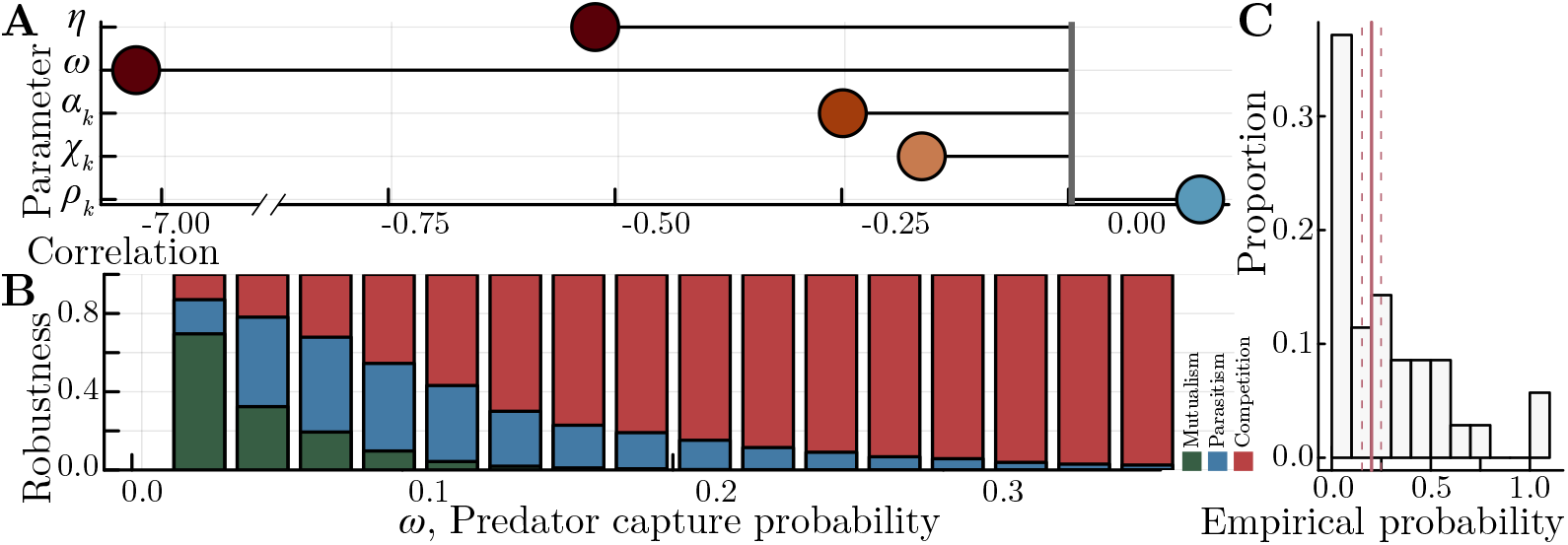
Parameter sensitivities and robustness of interaction modes. **A**. Correlations between parameters and the probability of apparent mutualism. Negative coefficients indicate smaller values of the parameter increase the probability that a random stable community described by Eq. 5 will exhibit apparent mutualism. **B**. Robustness, ℛ, of each interaction mode (Eq. 9) as a function of predator capture success probability, *ω*. Bins show the fraction of 10^6^ random stable communities described by Eq. 5 that result in competition, parasitism, or mutualism as capture probability, *ω*, is varied. Parameter definitions and ranges for random communities are defined in Table 1. **C**. Empirical predator capture probabilities from *N* = 8 published laboratory and field studies (35 experimental animal groups total; see *Robustness of apparent mutualism*). Pink lines show median = 0.2 (± 1 standard error measurement) capture success.

Given the outsized role of predator capture success, *ω*, in the emergence of apparent mutualisms (Fig. 3A, note *x*-axis scale), we further analyzed the robustness, ℛ (Eq. 9), of each apparent interaction type as capture success is systematically altered while all other parameters are drawn randomly from uniform distributions (Table 1; Fig. 3B). These quantities estimate the probability that a random stable community will result in apparent competition, parasitism, or mutualism as *ω* varies. At low *ω* values, the system predominantly exhibits mutualism, while higher *ω* values shift interactions towards competition (Fig. 3B). Intermediate *ω* values display a mix of modes, including parasitism, indicating that predator capture probability is the most important parameter for determining interaction mode prevalence. Data on capture success probabilities for predatory animals are limited. However, 35 empirical estimates across 8 studies of flying and swimming predators (Sweatman 1984; Moyer 1987; Norton 1991; Hirvonen & Ranta 1996; Sancho 2000; Cresswell & Quinn 2010; Corcoran & Conner 2016; Thiebault *et al*. 2016) indicate that capture probabilities are commonly within the range of values predicted by our model to lead to positive indirect interactions (median capture success of 0.2; Fig. 3C).

## DISCUSSION

Behavioral plasticity not only shapes direct species interactions but also fundamentally alters indirect interactions in ecological networks (Abrams 1987; Abrams 2010; Fahimipour & Anderson 2015). Our study shows that when prey adopt a common antipredator response, hiding (Lima & Dill 1990; Sih *et al*. 2004; Heithaus *et al*. 2009; Herberholz & Marquart 2012), two critical effects emerge. First, hiding reduces prey mortality by lowering exposure to predators. Second, it weakens density dependence by constantly shifting some fraction of the population in and out of a state that contributes little to intraspecific competition. Together, these effects reverse the traditional negative impact of predation: reduced hunting success leads to lower prey mortality and diminished competition, allowing both prey and predator biomass to increase under certain conditions (Figs. 2A–C).

Our results (*e*.*g*., Fig. 2) align with previous findings, that behaviors like hiding weaken interaction strengths and hence promote species coexistence. This pattern appears in models of prey shelter-seeking (Sih 1987) and refuge patch use in metacommunities (Křivan 1998). Positive indirect effects between apparent competitors also emerge in models with type III (Holling 1959; Murdoch 1969) consumer functional responses and in those where prey mortality scales with foraging time (Abrams 1987). More broadly, our model along with these prior insights suggest that behaviors weakening density regulation in response to encounters with other species ought to be key drivers of shifts in interaction valence. Predator-elicited hiding is one example, but similar effects may arise in diverse situations, for instance from trophically transmitted parasites that alter consumer foraging behavior after encounters with infected prey (Buck & Ripple 2017). This mechanism also bears a conceptual resemblance to the *hydra effect*, in which increased *per capita* mortality leads to higher population abundance (Abrams 2009). In both cases, interaction strength is modulated by a reduction in intraspecific competition. While our model involves a slightly different mechanism, the underlying logic is similar. Whether apparent mutualism might arise under hydra-like dynamics is an intriguing possibility for future work.

This dynamic may be particularly relevant in systems where predators exhibit low capture success, a common phenomenon in pursuit-and-escape interactions (Martin *et al*. 2022). While highly efficient predators exist (Fig.3C; Spencer *et al*. 2016), empirical data show that many taxa, including fish, birds, dragonflies, bats, and amphibians, operate with prey capture probabilities ranging from 1% to 20% (Moyer 1987; Sancho 2000; Sweatman 1984; Parrish 1993). Even among relatively successful predators, capture rates rarely exceed 50% (Cresswell & Quinn 2010; Corcoran & Conner 2016; Cresswell, Lind, *et al*. 2010; Thiebault *et al*. 2016; Hirvonen & Ranta 1996; Norton 1991) (Fig.3C). These values are consistent with the conditions under which our model predicts the emergence of positive indirect interactions. In practice, documenting failed predator attacks, behavioral state shifts, and changing indirect interactions in the field remains a significant challenge requiring detailed interaction time series from individual wild animals. Continued advances in machine vision (M. A. Gil & Hein 2017; Fahimipour, M. A. Gil, *et al*. 2023), drone technology (Koger *et al*. 2023), and continuous monitoring (Davies & Asner 2014; M. A. Gil, Michel, *et al*. 2025) provide promising tools for overcoming these obstacles.

Our approach can be readily extended to other behaviors or behaviorally mediated interactions, allowing for the incorporation of additional forms of organismal plasticity into population dynamics models. Inspired by classical coarse-grained models like reaction kinetics from chemistry (Michaelis, Menten, *et al*. 1913) and the functional response from ecology (Holling 1959; Real 1977; Auger & Roussarie 1994; Dawes & Souza 2013), we start by constructing a full system that includes subdivided populations that represent discrete behavioral or functional states in one or more species (Fig. 1). A reduced model can then be derived by assuming a time scale separation — solving for the fast variables, considering these auxiliary variables, and substituting them back into the slow manifold (Real 1977; Keener & Keizer 2002; O’Dwyer 2018; Holling 1959; Hein & Martin 2020; Muscarella & O’Dwyer 2020). The power of this, and other, coarse-graining methods (T. Gross & Feudel 2009) is that they can provide rapid analytical insights into which individual-level organismal behaviors might (or might not) influence dynamics at the scale of the entire ecosystem, or even when different microscopic animal behaviors map to the same macroscopic description of system dynamics (Wolpert *et al*. 2014; O’Dwyer 2018; D’Andrea *et al*. 2020).

While coarse-graining and time scale separation offer effective means to simplify complex systems, our findings reveal something perhaps more critical: routine animal behaviors can generate indirect effects that may defy intuition at the ecological network scale, such as predators benefiting from higher failure rates due to interactions within their network motif, or competitors transitioning into mutualists. These results show how even small behavioral changes can spread through ecological networks, leading to cascading effects that reshape community structure and function. Understanding such nontrivial relationships between scales is crucial for predicting and managing ecosystem resilience in a changing world, and may open the way for targeted interventions that modify specific organismal behaviors with known impacts on ecosystems.

## ACKNOWLEDGMENTS

We thank T. Gross for a lecture that inspired this work, B.T. Martin for help with data sources. A.K.F., M.A.G, and A.M.H. were supported by National Science Foundation grant no. EF-2222478. A.M.H. was supported by National Science Foundation grant nos. IOS-1855956 and IOS-2338596.

## REFERENCES

1. Abrams, P. A. Foraging time optimization and interactions in food webs. The American Naturalist 124, 80–96 (1984).

2. Abrams, P. A. Implications of flexible foraging for interspecific interactions: lessons from simple models. Functional Ecology, 7–17 (2010).

3. Abrams, P. A. Indirect interactions between species that share a predator: varieties of indirect effects. Predation: direct and indirect impacts on aquatic communities, 38–54 (1987).

4. Abrams, P. A. When does greater mortality increase population size? The long history and diverse mechanisms underlying the hydra effect. Ecology Letters 12, 462–474 (2009).

5. Abrams, P. A., Menge, B. A., Mittelbach, G. G., Spiller, D. A. & Yodzis, P. in Food webs: Integration of patterns & dynamics 371–395 (Springer, 1996).

6. Aufderheide, H., Rudolf, L., Gross, T. & Lafferty, K. D. How to predict community responses to perturbations in the face of imperfect knowledge and network complexity. Proceedings of the Royal Society B: Biological Sciences 280, 20132355 (2013).

7. Auger, P. M. & Roussarie, R. Complex ecological models with simple dynamics: From individuals to populations. Acta Biotheoretica 42, 111–136 (1994).

8. Barkai, A. & McQuaid, C. Predator-prey role reversal in a marine benthic ecosystem. Science 242, 62–64 (1988).

9. Beckerman, A. P., Uriarte, M. & Schmitz, O. J. Experimental evidence for a behavior-mediated trophic cascade in a terrestrial food chain. Proceedings of the National Academy of Sciences 94, 10735–10738 (1997).

10. Bender, E. A., Case, T. J. & Gilpin, M. E. Perturbation experiments in community ecology: theory and practice. Ecology 65, 1–13 (1984).

11. Bertness, M. D. & Callaway, R. Positive interactions in communities. Trends in Ecology & Evolution 9, 191–193 (1994).

12. Bshary, R., Grutter, A. S., Willener, A. S. & Leimar, O. Pairs of cooperating cleaner fish provide better service quality than singletons. Nature 455, 964–966 (2008).

13. Buck, J. C. & Ripple, W. J. Infectious agents trigger trophic cascades. Trends in Ecology & Evolution 32, 681–694 (2017).

14. Collins, J. P. & Cheek, J. E. Effect of food and density on development of typical and cannibalistic salamander larvae in Ambystoma tigrinum nebulosum. American Zoologist 23, 77–84 (1983).

15. Corcoran, A. J. & Conner, W. E. How moths escape bats: predicting outcomes of predator– prey interactions. Journal of Experimental Biology 219, 2704–2715 (2016).

16. Cresswell, W., Lind, J. & Quinn, J. L. Predator-hunting success and prey vulnerability: quantifying the spatial scale over which lethal and non-lethal effects of predation occur. Journal of Animal Ecology 79, 556–562 (2010).

17. Cresswell, W. & Quinn, J. L. Attack frequency, attack success and choice of prey group size for two predators with contrasting hunting strategies. Animal Behaviour 80, 643–648 (2010).

18. D’Andrea, R., Gibbs, T. & O’Dwyer, J. P. Emergent neutrality in consumer-resource dynamics. PLoS Computational Biology 16, e1008102 (2020).

19. Davies, A. B. & Asner, G. P. Advances in animal ecology from 3D-LiDAR ecosystem mapping. Trends in Ecology & Evolution 29, 681–691 (2014).

20. Dawes, J. & Souza, M. A derivation of Holling’s type I, II and III functional responses in predator–prey systems. Journal of Theoretical Biology 327, 11–22 (2013).

21. Dunne, J. A. & Williams, R. J. Cascading extinctions and community collapse in model food webs. Philosophical Transactions of the Royal Society B: Biological Sciences 364, 1711–1723 (2009).

22. Endler, J. A. Frequency-dependent predation, crypsis and aposematic coloration. Philosophical Transactions of the Royal Society of London. B, Biological Sciences 319, 505–523 (1988).

23. Fahimipour, A. K. & Anderson, K. E. Colonisation rate and adaptive foraging control the emergence of trophic cascades. Ecology Letters 18, 826–833 (2015).

24. Fahimipour, A. K., Anderson, K. E. & Williams, R. J. Compensation masks trophic cascades in complex food webs. Theoretical Ecology 10, 245–253 (2017).

25. Fahimipour, A. K., Gil, M. A., et al. Wild animals suppress the spread of socially transmitted misinformation. Proceedings of the National Academy of Sciences 120, e2215428120 (2023).

26. Gaiarsa, M. P., Kremen, C. & Ponisio, L. C. Pollinator interaction flexibility across scales affects patch colonization and occupancy. Nature Ecology & Evolution 5, 787–793 (2021).

27. Giese, A. C. Cannibalism and gigantism in Blepharisma. Transactions of the American Microscopical Society 57, 245–255 (1938).

28. Gil, M., Baskett, M. & Schreiber, S. Social information drives ecological outcomes among competing species. Ecology 100, e02835 (2019).

29. Gil, M. A. & Hein, A. M. Social interactions among grazing reef fish drive material flux in a coral reef ecosystem. Proceedings of the National Academy of Sciences 114, 4703–4708 (2017).

30. Gil, M. A., Michel, C. J., Olivetti, S., Sridharan, V. & Hein, A. M. Integrating Landscapes of Fear and Energy Reveals the Behavioural Strategies That Shape Predator–Prey Interactions. Ecology Letters 28, e70068 (2025).

31. Gilbert, J. J. Induction and ecological significance of gigantism in the rotifer Asplanchna sieboldi. Science 181, 63–66 (1973).

32. Gross, T. & Feudel, U. Analytical search for bifurcation surfaces in parameter space. Physica D: Nonlinear Phenomena 195, 292–302 (2004).

33. Gross, T. & Feudel, U. Local dynamical equivalence of certain food webs. Ocean Dynamics 59, 417–427 (2009).

34. Gurney, W. & Nisbet, R. M. Ecological dynamics (Oxford University Press, 1998).

35. Hastings, A. & Gross, L. J. Encyclopedia of Theoretical Ecology 4 (Univ of California Press, 2012).

36. Hein, A. M. & Martin, B. T. Information limitation and the dynamics of coupled ecological systems. Nature Ecology & Evolution 4, 82–90 (2020).

37. Heithaus, M. R., Wirsing, A. J., Burkholder, D., Thomson, J. & Dill, L. M. Towards a predictive framework for predator risk effects: the interaction of landscape features and prey escape tactics. Journal of Animal Ecology 78, 556–562 (2009).

38. Herberholz, J. & Marquart, G. D. Decision making and behavioral choice during predator avoidance. Frontiers in Neuroscience 6, 125 (2012).

39. Hirvonen, H. & Ranta, E. Prey to predator size ratio influences foraging efficiency of larval Aeshna juncea dragonflies. Oecologia 106, 407–415 (1996).

40. Holling, C. S. The components of predation as revealed by a study of small-mammal predation of the European Pine Sawfly. The Canadian Entomologist 91, 293–320 (1959).

41. Holt, R. D. Predation, apparent competition, and the structure of prey communities. Theoretical Population Biology 12, 197–229 (1977).

42. Holt, R. D. & Bonsall, M. B. Apparent competition. Annual Review of Ecology, Evolution, and Systematics 48, 447–471 (2017).

43. Huang, C. & Sih, A. Experimental studies on behaviorally mediated, indirect interactions through a shared predator. Ecology 71, 1515–1522 (1990).

44. Keener, J. P. & Keizer, J. E. in Computational Cell Biology 77–100 (Springer, 2002).

45. Khalil, H. K. Control of nonlinear systems (Prentice Hall, New York, NY, 2002).

46. Koger, B. et al. Quantifying the movement, behaviour and environmental context of group-living animals using drones and computer vision. Journal of Animal Ecology 92, 1357–1371 (2023).

47. Krivan, V. Effects of optimal antipredator behavior of prey on predator–prey dynamics: the role of refuges. Theoretical Population Biology 53, 131–142 (1998).

48. Kuznetsov, Y. A. Elements of applied bifurcation theory (Springer, 1998).

49. Lawton, P., Fahimipour, A. K. & Anderson, K. E. Interspecific dispersal constraints suppress pattern formation in metacommunities. Philosophical Transactions B 379, 20230136 (2024).

50. Lima, S. L. Stress and decision making under the risk of predation: recent developments from behavioral, reproductive, and ecological perspectives. Advances in the Study of Behavior 27, 215–290 (1998).

51. Lima, S. L. & Dill, L. M. Behavioral decisions made under the risk of predation: a review and prospectus. Canadian Journal of Zoology 68, 619–640 (1990).

52. Martin, B. T., Gil, M. A., Fahimipour, A. K. & Hein, A. M. Informational constraints on predator–prey interactions. Oikos 2022, e08143 (2022).

53. Martín, J. & López, P. Hiding time in refuge. Escaping from predators: an integrative view of escape decisions, 227e262 (2015).

54. Martín, J., López, P. & Cooper Jr, W. E. When to come out from a refuge: balancing predation risk and foraging opportunities in an alpine lizard. Ethology 109, 77–87 (2003).

55. Michaelis, L., Menten, M. L., et al. Die kinetik der invertinwirkung. Biochem. Z 49, 352 (1913).

56. Moyer, J. T. Quantitative observations of predation during spawning rushes of the labrid fish Thalassoma cupido at Miyake-jima, Japan. Japanese Journal of Ichthyology 34, 76–81 (1987).

57. Murdoch, W. W. Switching in general predators: experiments on predator specificity and stability of prey populations. Ecological Monographs 39, 335–354 (1969).

58. Muscarella, M. E. & O’Dwyer, J. P. Species dynamics and interactions via metabolically informed consumer-resource models. Theoretical Ecology 13, 503–518 (2020).

59. Nadell, C. D., Drescher, K. & Foster, K. R. Spatial structure, cooperation and competition in biofilms. Nature Reviews Microbiology 14, 589–600 (2016).

60. Nakajima, H. Sensitivity and stability of flow networks. Ecological Modelling 62, 123–133 (1992).

61. Neutel, A. M. et al. Reconciling complexity with stability in naturally assembling food webs. Nature 449, 599–602 (2007).

62. Norton, S. F. Capture success and diet of cottid fishes: the role of predator morphology and attack kinematics. Ecology 72, 1807–1819 (1991).

63. Novak, M. et al. Characterizing species interactions to understand press perturbations: what is the community matrix? Annual Review of Ecology, Evolution, and Systematics 47, 409–432 (2016).

64. O’Dwyer, J. P. Whence Lotka-Volterra? Conservation laws and integrable systems in ecology. Theoretical Ecology 11, 441–452 (2018).

65. Parrish, J. K. Comparison of the hunting behavior of four piscine predators attacking schooling prey. Ethology 95, 233–246 (1993).

66. Real, L. A. The kinetics of functional response. The American Naturalist 111, 289–300 (1977).

67. Sancho, G. Predatory behaviors of Caranx melampygus (Carangidae) feeding on spawning reef fishes: a novel ambushing strategy. Bulletin of Marine Science 66, 487–496 (2000).

68. Schmitz, O. J., Beckerman, A. P. & O’Brien, K. M. Behaviorally mediated trophic cascades: effects of predation risk on food web interactions. Ecology 78, 1388–1399 (1997).

69. Schmitz, O. J., Krivan, V. & Ovadia, O. Trophic cascades: the primacy of trait-mediated indirect interactions. Ecology Letters 7, 153–163 (2004).

70. Schmitz, O. J. & Suttle, K. B. Effects of top predator species on direct and indirect interactions in a food web. Ecology 82, 2072–2081 (2001).

71. Schreiber, S. J., Bürger, R. & Bolnick, D. I. The community effects of phenotypic and genetic variation within a predator population. Ecology 92, 1582–1593 (2011).

72. Schreiber, S. J. & Krivan, V. Holt (1977) and apparent competition. Theoretical Population Biology 133, 17–18 (2020).

73. Sih, A. Prey refuges and predator-prey stability. Theoretical Population Biology 31, 1–12 (1987).

74. Sih, A., Bell, A. M., Johnson, J. C. & Ziemba, R. E. Behavioral syndromes: an integrative overview. The Quarterly Review of Biology 79, 241–277 (2004).

75. Spencer, R. J., Van Dyke, J. U. & Thompson, M. B. The ethological trap: functional and numerical responses of highly efficient invasive predators driving prey extinctions. Ecological Applications 26, 1969–1983 (2016).

76. Sweatman, H. P. A field study of the predatory behavior and feeding rate of a piscivorous coral reef fish, the lizardfish Synodus englemani. Copeia, 187–194 (1984).

77. Thiebault, A., Semeria, M., Lett, C. & Tremblay, Y. How to capture fish in a school? Effect of successive predator attacks on seabird feeding success. Journal of Animal Ecology 85, 157–167 (2016).

78. Werner, E. E., Gilliam, J. F., Hall, D. J. & Mittelbach, G. G. An experimental test of the effects of predation risk on habitat use in fish. Ecology 64, 1540–1548 (1983).

79. White, J. W., Rassweiler, A., Samhouri, J. F., Stier, A. C. & White, C. Ecologists should not use statistical significance tests to interpret simulation model results. Oikos 123, 385–388 (2014).

80. Winnie Jr, J. A. Predation risk, elk, and aspen: tests of a behaviorally mediated trophic cascade in the Greater Yellowstone Ecosystem. Ecology 93, 2600–2614 (2012).

81. Wolpert, D. H., Grochow, J. A., Libby, E. & DeDeo, S. Optimal high-level descriptions of dynamical systems. arXiv preprint 1409.7403 (2014).

82. Yodzis, P. Diffuse effects in food webs. Ecology 81, 261–266 (2000).

