## Appendix 1 for "Behavioral plasticity and the valence of indirect interactions"

### *Appendix S1*: Behavioral plasticity and the valence of indirect interactions

Ashkaan K. Fahimipour[1,2,⋆], Michael A. Gil[3], and Andrew M. Hein[4]

[1] *Department of Biological Sciences, Florida Atlantic University, Boca Raton, FL, USA*
[2] *Center for Complex Systems, Florida Atlantic University, Boca Raton, FL, USA*
[3] *Department of Ecology and Evolutionary Biology, University of Colorado, Boulder, CO, USA*
[4] *Department of Computational Biology, Cornell University, Ithaca, NY, USA*

#### Density-dependent regulation in hiding prey

We analyze the sensitivity of our results to the assumption that hiding prey do not contribute to density dependence. To assess the robustness of our findings, we introduce an alternative model that relaxes this assumption by explicitly incorporating density dependence in hiding prey.

##### Modified model with density-dependent hiding prey

To test the impact of this assumption, we modify the prey growth function to include an additional term accounting for density dependence among hiders, where differences from the model in the *Main Text* are marked in red:

$$\dot{F}_k = r_k F_k \left(1 - \frac{F_k + \theta_k H_k}{K_k}\right) + x_k H_k - a_k F_k P, \tag{1a}$$

$$\dot{H}_k = \tilde{\omega} a_k F_k P - x_k H_k, \tag{1b}$$

$$\dot{P} = \omega \sum_k e_k a_k F_k P - mP. \tag{1c}$$

Here, $\theta_k \in [0, 1]$ determines the extent to which hiding prey contribute to density dependence, with all other parameters as in the *Main Text*. Setting $\theta_k = 0$ recovers our original model (see Eq. 1 in the *Main Text*), while $\theta_k = 1$ corresponds to the case where hiders and foragers contribute equally to intraspecific competition. Since the nondimensionalization and time scale separation procedure remains unchanged, the resulting model is similar to Eq. 5 in the *Main Text*. Applying the same substitutions as before, we obtain the nondimensionalized system:

$$\dot{f}_1 = \rho_1 f_1 (1 - f_1 - \theta_1 h_1) + \chi_1 h_1 - \alpha_1 f_1 p, \tag{2a}$$

$$\dot{h}_1 = \tilde{\omega} \alpha_1 f_1 p - \chi_1 h_1, \tag{2b}$$

$$\dot{f}_2 = \rho_2 f_2 (1 - f_2 - \theta_2 h_2) + \chi_2 h_2 - \alpha_2 f_2 p, \tag{2c}$$

$$\dot{h}_2 = \tilde{\omega} \alpha_2 f_2 p - \chi_2 h_2, \tag{2d}$$

$$\dot{p} = \omega p (\alpha_1 f_1 + \eta \alpha_2 f_2) - p. \tag{2e}$$

The steady-state values of $h_k^\star$ are found by setting $\dot{h}_k(h_k^\star) = 0$ in Eq. 2. Substituting $h_k^\star$ back into the predator and total prey equations yields a model structurally identical to that in the *Main Text*, so we omit it here for brevity. However, the additional term modifies the prey growth function $\phi_k(n_k, p)$, now given by

$$\phi_k(n_k, p) = \frac{\chi_k(1 - n_k) + \tilde{\omega}\alpha_k p(1 - \theta_k n_k)}{(\chi_k + \tilde{\omega}\alpha_k p)^2}. \tag{3}$$

This formulation reveals that the positive effect of hiding ($\tilde{\omega}\alpha_k p$) is now attenuated by prey themselves, according to the factor $(1 - \theta_k n_k)$.

#### Analysis of press perturbations

The additional complexity of the model permits an analysis of press perturbations via simulation. The numerical results should correspond proportionally to the matrix $\mathbf{M} = -\mathbf{J}^{-1}$ derived in the *Main Text* (Aufderheide *et al.* 2013). Specifically, we conducted numerical simulations to investigate the effects of permanent press perturbations on species abundances. The system was first integrated to steady state under baseline conditions. At that point, we applied a permanent 10% increase to the intrinsic growth rate ($\rho_k$) of a single prey species while holding all other parameters constant. The system was then integrated forward until it reached a new steady state (Fig. S1A).

To quantify species responses to press perturbations (*i.e.*, indirect effects), we computed the log response ratio for each prey species $k \in [1, 2]$, defined as

$$\mathcal{L}_k = \log \frac{n_{\text{perturbed}}}{n_{\text{baseline}}}, \tag{4}$$

where $n_{\text{perturbed}}$ and $n_{\text{baseline}}$ denote the equilibrium abundances after and before perturbation, respectively. This metric allowed us to assess how changes in one species' growth rate influenced the equilibrium structure of the system and valence of interactions between prey: *apparent competition* occurs when both prey negatively affect each other, $\max(\mathcal{L}_1,\ \mathcal{L}_2) < 0$; *apparent parasitism* occurs when one prey benefits at the expense of the other, $\min(\mathcal{L}_1,\ \mathcal{L}_2) < 0 < \max(\mathcal{L}_1,\ \mathcal{L}_2)$; and *apparent mutualism* occurs when both prey benefit from one another, $0 < \min(\mathcal{L}_1,\ \mathcal{L}_2)$. All simulations were performed using the LSODA implementation (Hindmarsh 1992) in *DifferentialEquations.jl* (Rackauckas & Nie 2017) within the Julia v1.11.2 programming language. The system was considered to have reached steady state when the maximum rate of change across all species fell below $10^{-7}$.

We begin by defining a parameter set and parameter ranges under which apparent mutualism occasionally arises (*see* Fig. S1 caption) and then examine how the frequency of apparent mutualism and parasitism shifts as the density-dependent parameter $\theta$ increases, while drawing other parameters randomly from relevant distributions (*see* Fig. S1 caption). The results, summarized in Fig. S1B, show minimal effects of increasing $\theta$ except at extreme values near $\theta = 1$, where density dependence is strongest.

This leads to two key insights. First, even when hiders contribute substantially to intraspecific competition, positive indirect effects can still emerge. Second, the fact that these effects disappear only in the extreme and biologically implausible case where hiders compete as strongly as foragers reinforces the idea that shifts in prey interaction valence are driven by predator-mediated changes in intraspecific competition. Ultimately, this highlights the central role of predation in shaping

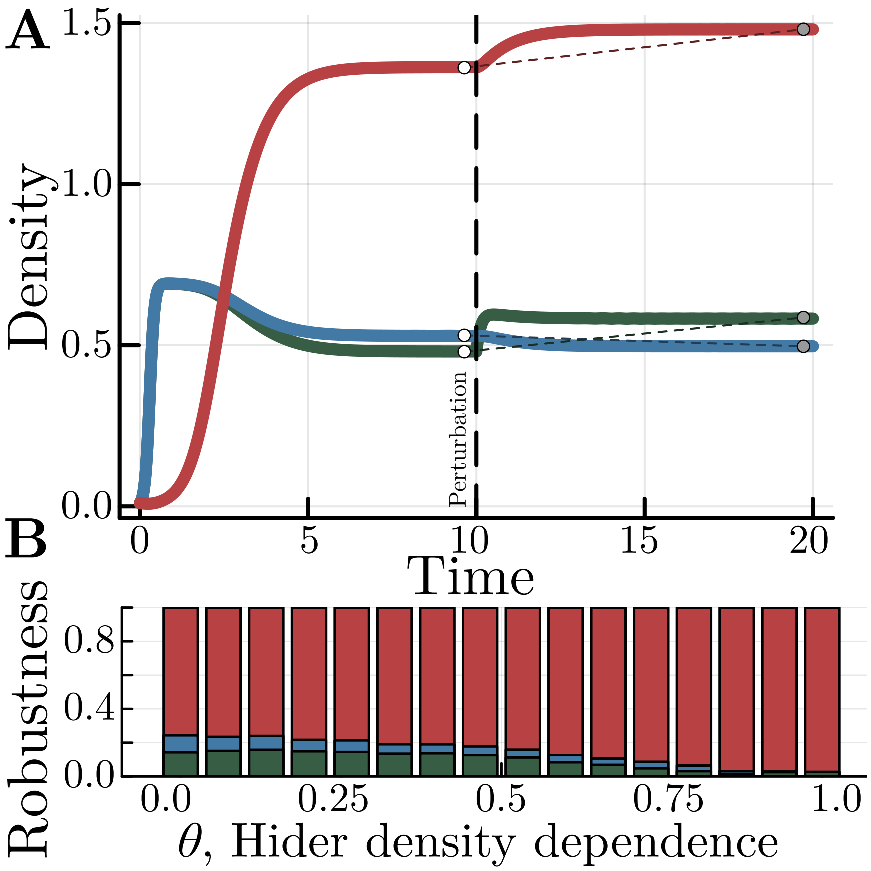


Supplementary Figure 1: **Simulation-based press perturbations. A**. Illustration of a single simulated press perturbation (red is the predator, green and blue are prey spp. 1 and 2 respectively). At time $t = 10$, the growth rate of prey sp. 1, $\rho_1$ is permanently increased by 10%. The values of $n_{\text{perturbed}}$ and $n_{\text{baseline}}$ for all species are marked by grey and white dots. **B**. Robustness of apparent mutualism as the strength of prey density dependence $\theta_k$ is increased. Parameters for the $10^5$ simulations are: $\rho_k = 14.5$, $\alpha_k \sim \mathcal{U}(5, 15)$, $\omega \sim \mathcal{U}(0.0, 0.25)$, $\eta = 1$, $\chi_k = \rho_k \cdot \mathcal{U}(1, 5)$. Colors for panel **B** are: red, competition; blue, parasitism; green, mutualism.

indirect prey interactions, even in systems with strong density dependence.

#### References

Aufderheide, H., Rudolf, L., Gross, T. & Lafferty, K. D. How to predict community responses to perturbations in the face of imperfect knowledge and network complexity. *Proceedings of the Royal Society B: Biological Sciences* **280,** 20132355 (2013).

Hindmarsh, A. *ODEPACK. A collection of ODE system solvers* tech. rep. (Lawrence Livermore National Lab, Livermore, CA (United States), 1992).

Rackauckas, C. & Nie, Q. DifferentialEquations.jl–a performant and feature-rich ecosystem for solving differential equations in Julia. *Journal of Open Research Software* **5** (2017).
